# Catalytic Bacterial Nanocellulose Composite That Captures and Degrades PET Microplastics

**DOI:** 10.64898/2026.06.05.730390

**Authors:** Brittany N. Pitt, Dayana Guillen, Allie C. Obermeyer

**Affiliations:** Department of Chemical Engineering, Columbia University, New York, NY 10027

**Keywords:** biocatalysis, enzyme immobilization, PET hydrolysis, bacterial nanocellulose, microplastic degradation

## Abstract

Microplastic (MP) pollution is a growing environmental concern, with wastewater treatment plants (WWTPs) serving as a critical yet insufficient barrier. Although WWTPs can remove up to 99% of MPs from influent streams, a large fraction of the sequestered MPs is concentrated into sewage sludge and subsequently redistributed to soil and waterways via biosolids. Here, we report a biosynthetic approach to address MP redistribution by coupling microplastic removal with catalytic depolymerization, using enzyme-functionalized bacterial nanocellulose (BNC) filtration materials. As a proof-of-concept, we fused a cellulose-binding domain (CBD) to the thermostable PET hydrolase HotPETase, creating a bifunctional protein, CBD-PETase, that binds to a BNC scaffold and degrades PET microplastic (µPET) captured within the scaffold. We showed that BNC-CBD-PETase composites sequestered irregularly shaped µPET within hierarchical pore networks and generated PET degradation products proportionally to enzyme loading. We further established a co-culture strategy, in which *Saccharomyces cerevisiae* was engineered to secrete CBD-PETase during BNC scaffold biosynthesis by *Komagataeibacter xylinus*, which enabled one-pot fabrication of an active BNC-CBD-PETase composite without separate protein purification. Notably, co-culture-derived BNC-CBD-PETase composites retained catalytic activity for at least 24 weeks under dry, ambient storage. This platform provides a route towards a scalable and renewable filtration material with the potential for integration across multiple treatment stages within WWTPs.

## Introduction

Microplastic (MP) pollution has emerged as a pervasive environmental challenge, with increasing detection across both aquatic and terrestrial ecosystems. Owing to their small size and heterogeneity, these particles are not only difficult to remove, but can also accumulate in living organisms, leading to persistent bioaccumulation.^1–3^

Wastewater is the dominant pathway by which MPs travel from urban sources and into the broader environment.^4,5^ This positions wastewater treatment plants (WWTPs) as a critical point for intervention. WWTPs remove MPs through a combination of physical, biological, and membrane-based treatment processes. The preliminary and primary treatment stages rely on screening, sedimentation, coagulation, and flocculation to remove larger suspended particles and coarse MPs (>500 µm), accounting for nearly 80% of overall microplastic removal.^5–9^ The secondary treatment stage employs microorganisms to degrade the dissolved and suspended organic matter within wastewater. Although MPs are not readily degraded during this stage, fibrous MPs are often adsorbed onto biomass surfaces and removed alongside sludge during flocculation and settling, accounting for the removal of 10-15% of MPs entering WWTPs.^5,10–12^ The tertiary treatment stage further removes finer particles (>20 µm) from water entering the inlets, using advanced filtration technologies such as rapid sand filtration (97% removal of MPs), membrane disc filters (50-98.5% removal of MPs), and membrane bioreactors (99.9% removal of MPs).^13^ Tertiary systems operate primarily through physical retention, trapping particles within porous surfaces without degrading them, thereby concentrating MPs within filter residues and sewage sludge.^14^ Although WWTPs remove up to 99% of MPs from their influent water streams, these particles are not degraded, but instead concentrated into sewage sludge, where >65% of MPs accumulate, and are subsequently redistributed into soils and waterways via biosolids.^5–12^ This points to a fundamental gap in current treatment approaches: capture is not coupled to degradation. Consequently, strategies that combine the removal of MPs with their degradation must be developed to prevent further accumulation of MPs in the environment.

Current efforts to reduce MPs released from WWTPs largely focus on improving removal from effluent streams through advanced tertiary filtration technologies, with comparatively little emphasis on preventing microplastic accumulation in the sludge product from primary and secondary treatment stages.^5,10,11,13^ We propose that interception strategies should also be integrated at these stages to capture microplastics both before they are concentrated in sludge, and after they are released into effluent streams. In this context, filtration materials with dynamic pore sizes that can both capture and degrade microplastics may offer a versatile, scalable approach for integration across multiple stages of wastewater treatment.

In one promising approach, bacterial nanocellulose (BNC) can be used to create such filtration materials. BNC is produced by the fermentation of a subset of bacteria, such as *Rhizobium, Agrobacterium, Pseudomonas, Komagataeibacter, Alcaligenes* and *Sarcina ventriculi.*^15^ It is secreted as pure cellulose nanofibers and assembled into a fibrillar network outside of the bacterial cells. Unlike plant-derived cellulose, it is free of lignin and hemicellulose, resulting in a highly pure and crystalline material.^16,17^ Importantly, the pore size and network density of BNC can be tuned from the nanoscale to several microns by altering the fermentation or adding post-processing steps, making it a highly adaptable filtration scaffold.^18–24^ These qualities position BNC as an exceptional scaffold for developing filtration materials for the removal of microplastics. Excitingly, BNC can also be engineered to incorporate both organic and inorganic materials to bestow additional properties,^22,25–29^ such as dye removal from wastewater facilitated by mesoporous polydopamine and palladium nanoparticles ^22^ and antibiotic degradation by the β-lactam hydrolyzing enzyme TEM1.^25^ To impart the inherent filtration ability of BNC with plastic degradation capacity, we sought to incorporate plastic-degrading enzymes into a BNC scaffold, thereby obtaining a composite filtration material that both captures and degrades MPs.

The enzymatic degradation of plastics has advanced considerably over the past decade, though progress has been uneven across polymer types. The first evidence of microbial plastic degradation dates to 1975, when *Flavobacterium* sp. was found to produce nylonase enzymes capable of hydrolyzing nylon oligomers in wastewater near a nylon factory.^30^ Since then, numerous plastic-degrading enzymes have been discovered.^31^ In 2012, the discovery of leaf-branch compost cutinase (LCC)^32^ introduced a thermostable enzyme scaffold capable of operating near the glass transition temperature of poly(ethylene terephthalate) (PET), making it particularly attractive for industrial PET depolymerization. The field accelerated further in 2016 with the discovery of the PET-hydrolyzing enzyme IsPETase,^33^ which demonstrated that microorganisms could naturally evolve the ability to depolymerize PET into the constituent monomers under ambient conditions. Following these discoveries, extensive protein engineering has produced improved IsPETase variants, including ThermoPETase,^34^ DuraPETase,^35^ HotPETase,^36^ and FAST-PETase.^37^ These variants have enhanced thermostability, catalytic efficiency, and substrate accessibility compared to the IsPETase predecessor. Parallel engineering of the LCC scaffold has also led to the development of LCC^ICCG^ ^38^ and TurboPETase,^39^ which also display significant improvement over the LCC predecessor.

Among currently engineered PETases, HotPETase is particularly well-suited for integration into BNC-based filtration systems. Developed in 2022 through rounds of directed evolution, HotPETase can depolymerize untreated post-consumer PET with crystallinities as high as 41% without pretreatment, making it one of the few enzymes capable of acting on the partially crystalline microplastics commonly found in environmental samples.^36,40^ In addition, the high operating temperature of HotPETase means that the enzyme remains largely inactive at the ambient conditions used to capture microplastics derived from PET (µPET). This separation of microplastic capture from degradation would minimize monomer release into the effluent during treatment; µPET degradation can therefore be activated on demand via controlled heating.

Herein, we report a versatile, fully biodegradable cellulose-enzyme composite that enables capture and catalytic depolymerization of µPET (Scheme 1). The combination of capture and degradation functionality into renewable-material platforms offers a practical route towards re-structuring WWTPs as barriers to microplastic pollution. By engineering a fusion between a thermally robust PET hydrolase, HotPETase,^36,41^ and a cellulose-binding domain (CBD),^42,43^ we enable localization of catalytic activity within a bacterial nanocellulose scaffold. This scaffold thus acts as a catalytic filter, enabling both sequestration and degradation of µPET. Mechanistically, we show that catalytic performance is limited by interfacial constraints arising from the inherent restricted accessibility of catalytic sites on semicrystalline µPET. This system therefore provides a foundation for material design, with the potential to improve as hydrolases are further optimized to overcome these limitations. Additionally, this system has the potential to be diversified through tuning mechanical properties such as pore size^18,23,44^ and tensile strength^45,46^ to meet specifications for WWTP filtration devices as well as by immobilizing different proteins to treat other wastewater microcontaminants as protein engineering efforts evolve.^20,47–49^ We further establish a co-culture strategy^25,26^ for in situ assembly of the composite during cellulose biosynthesis, providing a route towards scalable material fabrication.^20,28,50^

**Scheme 1.**
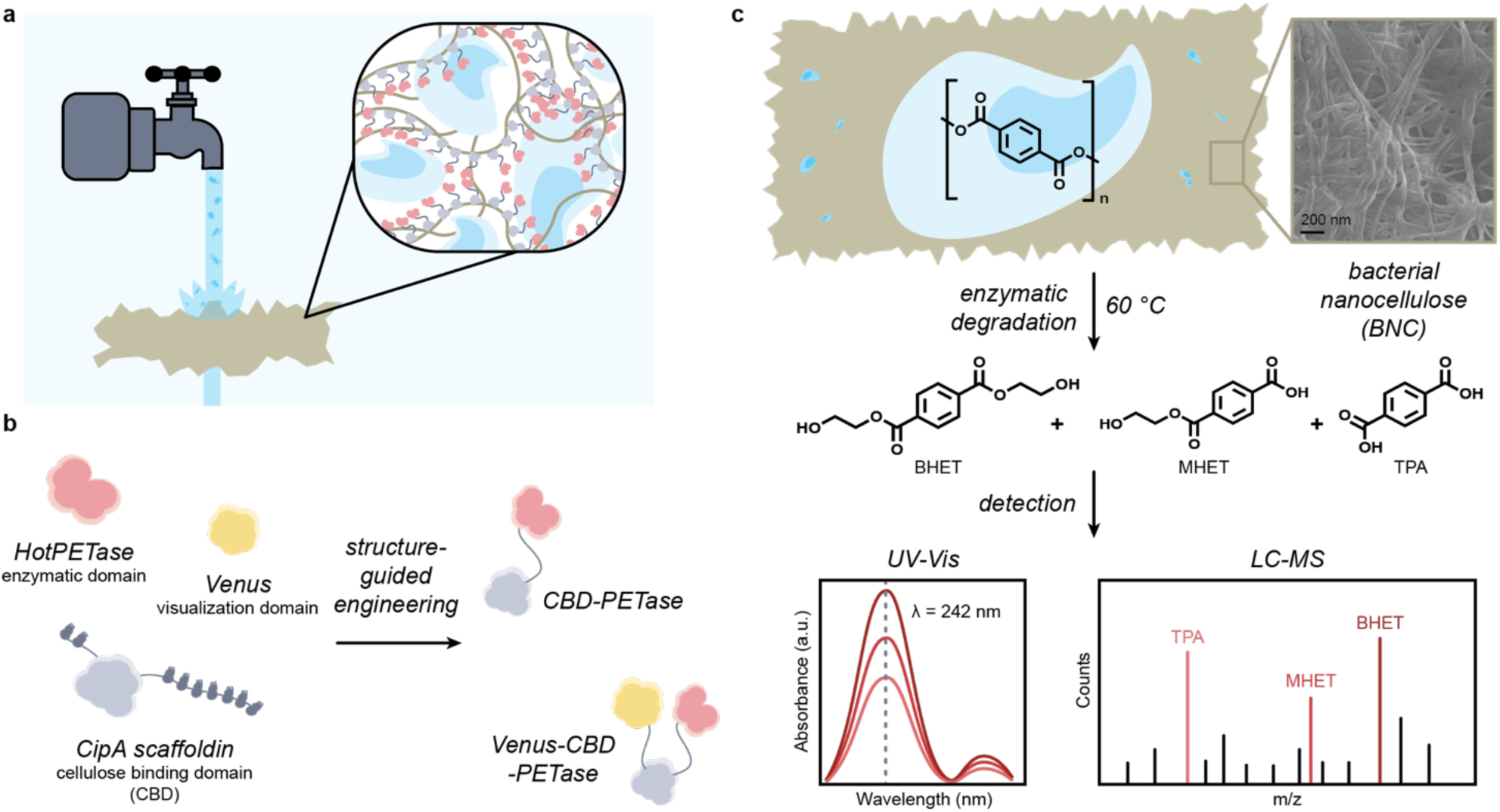
Overview of BNC-CBD-PETase Composite Fabrication and Function. (a) BNC-CBD-PETase composite directly sequesters μPET from water. (b) Structure-guided engineering is used to create a bi-functional protein that binds to cellulose, as well as degrades μPET. (c) Schematic of μPET capture in a functional BNC filter followed by enzymatic degradation, representative SEM of the BNC filter material, and schematic data of the degradation of μPET to the constituent monomers bis(2-hydroxyethyl) terephthalate (BHET), mono-(2-hydroxyethyl) terephthalic acid (MHET) and terephthalic acid (TPA) as monitored by UV-Vis absorbance and LC-MS analysis.

## Results and Discussion

To establish BNC functionalized with CBD-PETase (BNC-CBD-PETase) as an active scaffold for µPET capture and degradation (Scheme 1), we first verified that each component of the system retained functionality after engineering and assembly. First, rational engineering was used to modify the HotPETase enzyme such that it could be anchored to bacterial nanocellulose with sufficient flexibility to move within the composite and interact with sequestered microplastics. This was achieved by using the CipA cellulosome protein, from the bacterium *C. thermocellum*, as a basis for structure-guided engineering (Scheme 1b). The CipA cellulosome protein consists of a cellulose binding domain (CBM3a), preceded and followed by 2 and 7 cohesion domains, respectively. The cohesion domains interact with dockerin-containing enzymes secreted by the bacteria, forming a multi-enzyme complex, while CBM3a anchors this complex to the cellulose to be degraded. Consequently, the CipA scaffolding protein forms an enzyme complex that both binds to and degrades cellulose.^42,43^ To mimic the dual cellulose-binding and enzymatic functionality of CipA, we genetically fused the cellulose-binding domain CBM3a and the native linker domain to the N-terminus of the HotPETase enzyme. In addition, a flexible linker was added between the semi-rigid CBM3a linker and the HotPETase domain to minimize steric hindrance, creating the CBD-PETase enzyme discussed herein. This approach used the same principle as the CipA scaffolding protein to immobilize enzymes to cellulose but eliminated the multi-step assembly of dockerin-containing enzymes with the CipA scaffold. A fluorescent analog was also engineered to visualize the protein interactions with sequestered microplastics via fluorescence microscopy.

### Assessment of Thermal Stability and µPET Depolymerization of CBD-PETase Fusion

Given that PET degradation is most efficient near the glass transition temperature (T_g_) of the polymer, when ester bond cleavage sites are most accessible, the thermal stability of the engineered enzyme was characterized. The enzyme CBD-PETase was heated (1°C/min) from room temperature to 90 °C, and the thermal unfolding was monitored by changes in absorbance at 280 nm. Absorbance at this wavelength provides a composite readout of changes in the local environment of aromatic residues, as well as scattering associated with protein aggregation.

Analysis of the absorbance as a function of temperature for the CBD-PETase fusion enzyme shows two thermal transitions, as expected for a two-globular-domain protein. The first, occurring at ∼78 °C, likely corresponds to the unfolding of the CBM3a domain, and aligns with the melting temperature (T_m_) observed when differential scanning fluorimetry (DSF) was used to probe CBM3a-containing proteins,^51^ while the second, at ∼86 °C, likely corresponds to the unfolding of the HotPETase domain. A parallel experiment was conducted with the single-domain HotPETase enzyme as a control and, as expected, only a single transition was observed at ∼82 °C, consistent with the reported T_m_ ^36^ of HotPETase (Figure 1a). Consequently, we concluded that the CBD-PETase fusion does not undergo significant conformational changes at temperatures below 70 °C and therefore has the thermal stability to function near the T_g_ of the high-crystallinity µPET (SI Figure 2) used in this study.

**Figure 1.**
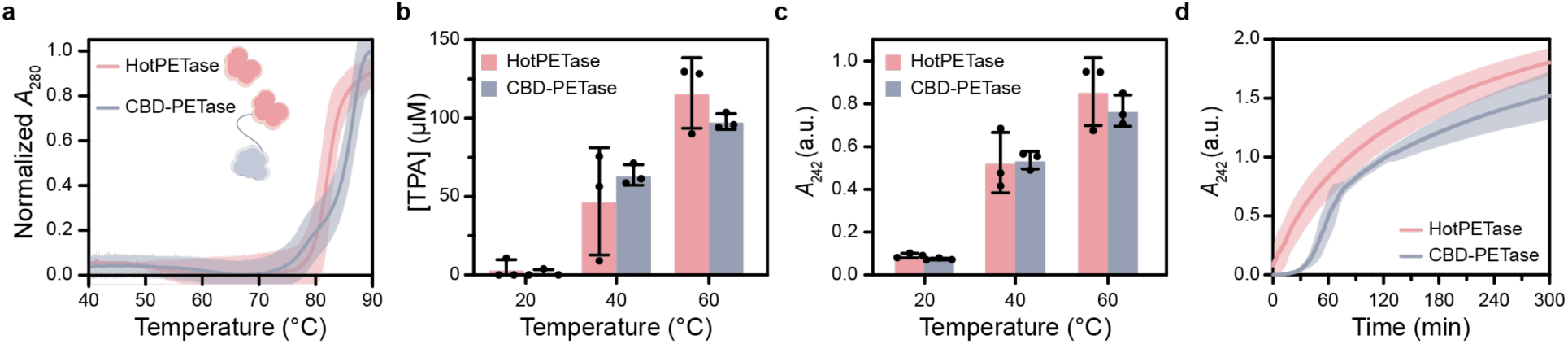
Assessment of Thermal Stability and μPET Depolymerization of CBD-PETase Fusion. (a) Plot of absorbance at 280 nm as a function of temperature for a solution of enzyme (0.45 μM) in 50 mM glycine buffer (Gly-OH), pH 9.2. Data were normalized to offset differences in absorbances resulting from differing amino acid composition. The temperature ramp rate was 1°C/min. Quantitative assessment of the concentration of (b) TPA as measured by LC-MS, (c) total soluble aromatic products as measured by absorbance at 242 nm, at 20 °C, 40 °C, and 60 °C. Individual data points from 3 independent replicates shown with average (bar) and standard deviation (error bars). (d) The production of soluble aromatic products was continuously monitored by absorbance measurement at 242 nm over 300 min for both the CBD-PETase fusion and HotPETase at 60 °C. Graphs (b), (c) and (d) were generated from experiments using 0.45 μM enzyme and 2 g L^-1^ of μPET in Gly-OH (pH 9.2). All line graphs correspond to the average of n = 2, except for CBD-PETase in graph (d), where n = 3 for accurate representation of the lag-phase claim. Shaded regions of all line graphs represent the standard deviation.

**Figure 2.**
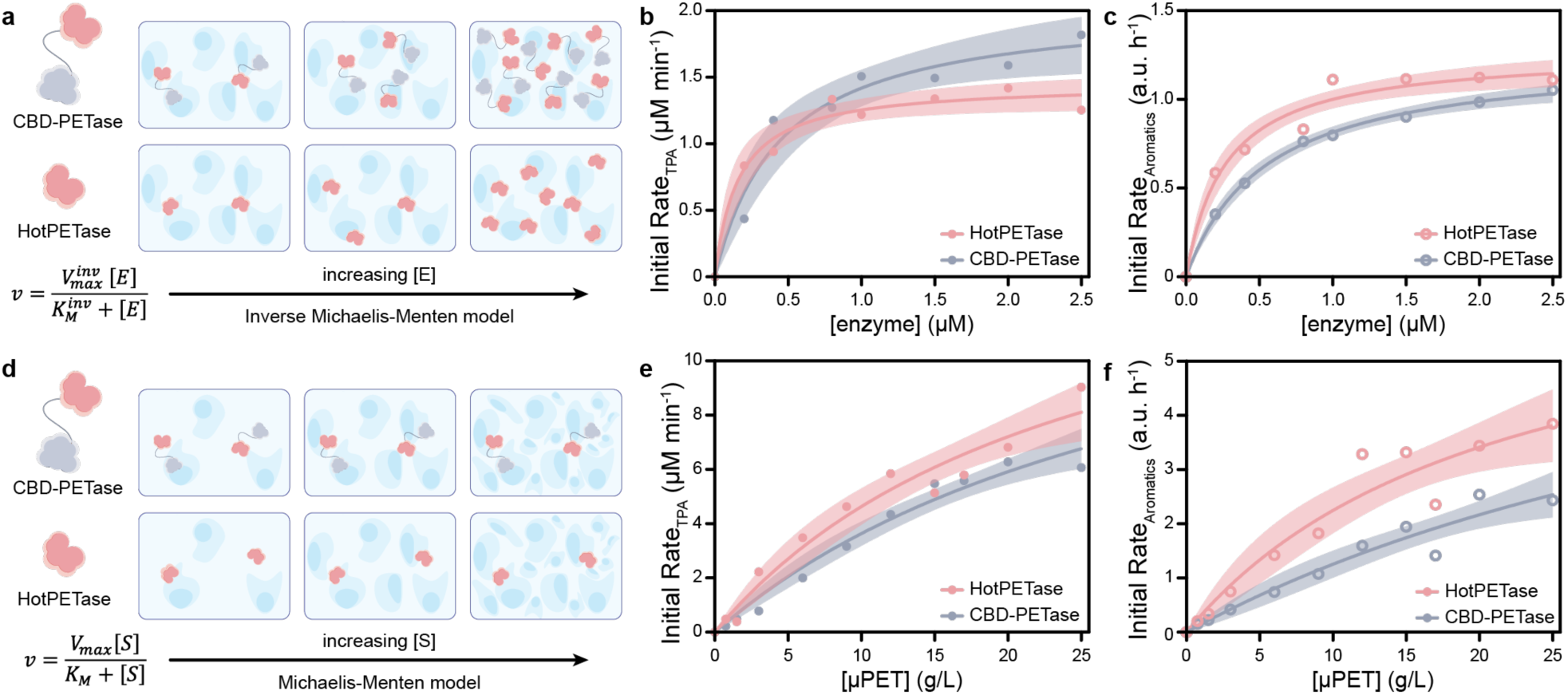
Inverse and Conventional Michaelis-Menten Models to Quantitatively Describe Enzymatic μPET Depolymerization in Free Solution. (a) Schematic depicting the enzyme-substrate relationship probed by the inverse Michaelis-Menten model. Inverse Michaelis-Menten fits as a function of enzyme concentration using initial production rate of (b) TPA, (c) soluble aromatics. (d) Schematic depicting the enzyme-substrate relationship probed by the conventional Michaelis-Menten model. Conventional Michaelis-Menten fits as a function of substrate concentration using initial production rate of (e) TPA, (f) soluble aromatics. Data points indicate the average of 3 independent experiments, the line shows the fit to the appropriate model, and the shaded region corresponds to the 95% CI of the fit.

Based on this thermal profile, enzymatic degradation assays were performed at 20, 40, and 60 °C to determine the optimal degradation temperature for the CBD-PETase fusion enzyme, relative to the original HotPETase. Using identical enzyme and substrate concentrations (0.45 µM enzyme, 2 g L^-1^ µPET), product formation was quantified after 3 h by LC-MS and UV-Vis absorbance for both HotPETase and CBD-PETase (Figure 1b,c). LC-MS quantification of the TPA degradation product indicated that increasing the temperature increased monomer production for both CBD-PETase and HotPETase (Figure 1b). Consistent with this trend, absorbance measurements at 242 nm (*A*_242_), a proxy for the concentration of total aromatic PET degradation products (SI Figure 3), also showed strong temperature dependence with maximal product formation at 60 °C. The increase in depolymerization of µPET with increasing temperature supports a surface site-limited degradation mechanism, characteristic of catalytic systems containing semicrystalline substrates such as PET, in which higher temperatures increase polymer flexibility and, consequently, improve the enzyme’s access to reactive sites.^52,53^ Critically, fusion of the enzyme to the CBM3a domain did not compromise the structural integrity nor the catalytic activity of the HotPETase domain.

**Figure 3.**
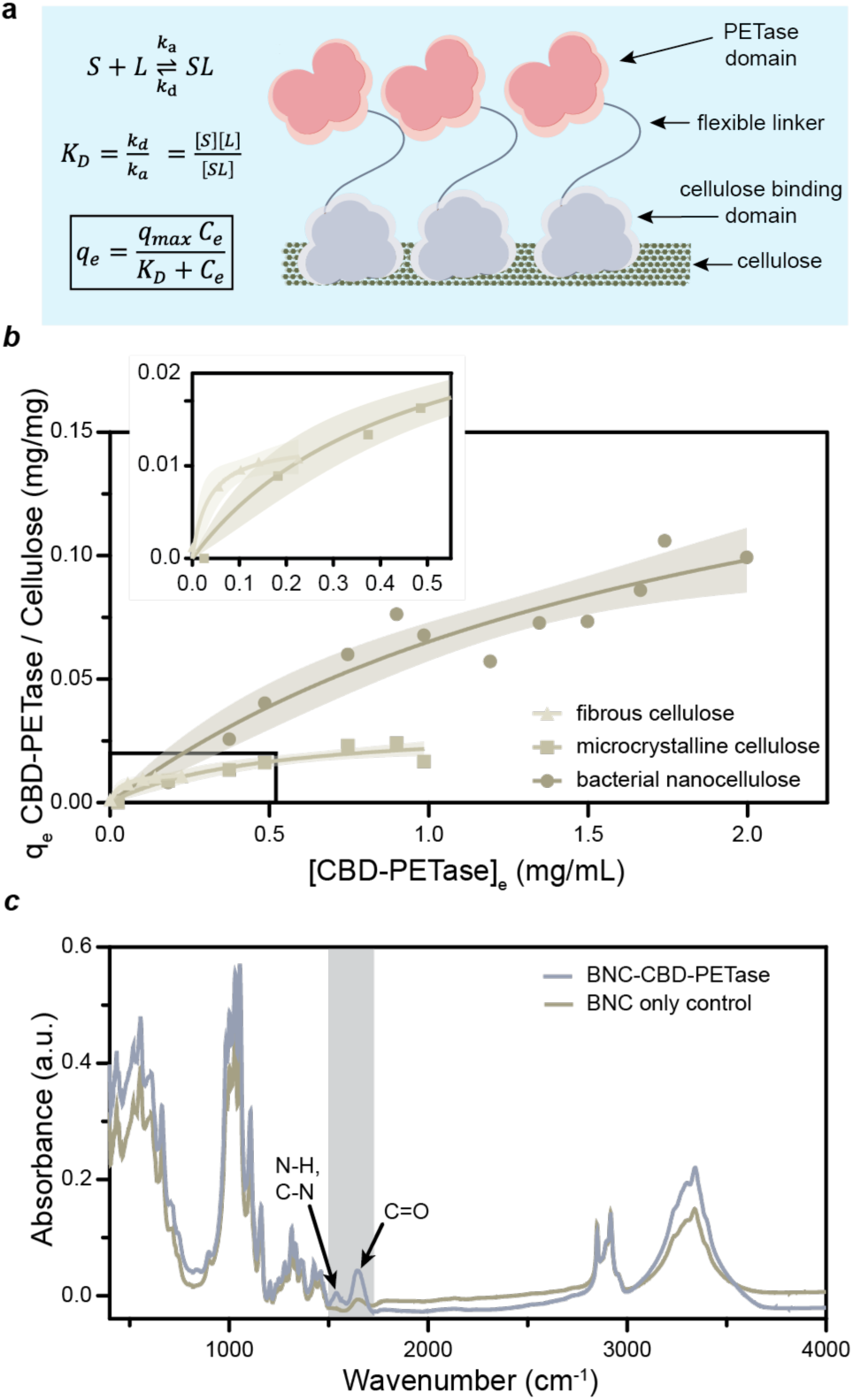
Langmuir Binding Model Describes the Immobilization of CBD-PETase to Cellulose. (a) Schematic illustrating the immobilization of CBD-PETase to cellulose. (b) Experimental data fitted to Langmuir binding isotherms describe the immobilization of CBDPETase to fibrous cellulose, microcrystalline cellulose, and bacterial nanocellulose. Data points represent the average of n=3 replicates, the line shows the fit to the Langmuir binding model, and the shaded region corresponds to the 95% confidence interval of the fit. (c) FT-IR data show efficient immobilization of CBD-PETase onto bacterial nanocellulose at low Ci (0.02 mg/mL) with new or higher intensity peaks in the amide region (1550-1770 cm-1).

Following thermal characterization, we monitored the depolymerization of µPET by the free enzymes to provide a basis for subsequent kinetic experiments. This was achieved through continuous quantification of the concentration of soluble aromatic PET degradation products generated by the enzymes via *A*_242_, as a function of time. Upon addition to the substrate, the CBD-PETase enzyme experienced a 20 min lag phase, followed by a linear increase in *A*_242_ for 1 h and then a plateau. On the other hand, a lag phase was not observed for the HotPETase enzyme, and a linear increase in *A*_242_ was observed for 1 h, followed by a plateau (Figure 1d). The observed lag phase for the fusion enzyme may partially arise from transient, non-productive interactions between the free CBM3a domain and µPET. It has been established that CBDs can bind to PET via π-stacking and polar interactions, likely reflecting the chemical similarities between the PET polymer and many carbohydrates.^54–63^ In the engineered CBD-PETase enzyme, there is both a semi-rigid and a flexible linker separating the CBM3a domain from the HotPETase domain. This could position some CBM3a-bound µPET far from the HotPETase catalytic site, reducing productive hydrolysis. In this case, we hypothesize that substrate competition between the two domains achieves equilibrium in ∼20 min, after which degradation proceeds steadily and linearly for an additional hour.

### Inverse and Conventional Michaelis-Menten Models Quantitatively Describe Enzymatic µPET Depolymerization

Kinetic characterization was performed by measuring the initial rate of product formation after incubation at 60 °C for 1 h as a function of increasing enzyme concentration (0–2.5 µM), with a fixed substrate concentration of 2 g L⁻¹. Quantification of the initial production rate of TPA and soluble aromatics revealed product saturation at enzyme concentrations exceeding sub-µM (Figures 2b, 2c). This was again consistent with a surface-limited catalytic process where, under fixed experimental conditions, the number of accessible reaction sites on the µPET surface is finite. Therefore, the inverse Michaelis-Menten framework, in which enzyme concentration serves as the saturating variable while the available surface sites are held constant (SI Derivation 1), was applied to the data to quantitatively describe the degradation kinetics (Table 1). Fitting the initial production rate of TPA yielded apparent 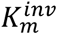values of 0.16 ± 0.04 µM and 0.44 ± 0.13 µM for HotPETase and CBD-PETase, respectively (Figure 2b). On the other hand, fitting the initial production rate of soluble aromatics yielded slightly higher apparent 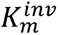of 0.27 ± 0.06 µM, and 0.54 ± 0.07 µM for HotPETase and CBD-PETase, respectively (Figure 2c).

**Table 1.** Kinetic Parameters for µPET Degradation.

|  | Production of Soluble Aromatics |  |  |  |  |
| --- | --- | --- | --- | --- | --- |
| | $V_{max}^{inv}$ (a.u. h<sup>-1</sup>) | $K_m^{inv}$ ($\mu\text{M}$) | $V_{max}$ (a.u. h<sup>-1</sup>) | $K_m$ (g L<sup>-1</sup>) | $\Gamma_{max}^{kin}$ |
| HotPETase | $1.27 \pm 0.06$ | $0.27 \pm 0.06$ | $7 \pm 2$ | $22 \pm 12$ | 0.04 |
| CBD-PETase | $1.26 \pm 0.05$ | $0.54 \pm 0.07$ | $8 \pm 5$ | $50 \pm 40^\dagger$ | 0.03 |
|  | Production of TPA |  |  |  |  |
| | $V_{max}^{inv}$ ($\mu\text{M min}^{-1}$) | $K_m^{inv}$ ($\mu\text{M}$) | $V_{max}$ ($\mu\text{M min}^{-1}$) | $K_m$ (g L<sup>-1</sup>) | $\Gamma_{max}^{kin}$ |
| HotPETase | $1.45 \pm 0.07$ | $0.16 \pm 0.04$ | $16 \pm 4$ | $25 \pm 11$ | 0.02 |
| CBD-PETase | $2.05 \pm 0.18$ | $0.44 \pm 0.13$ | $16 \pm 4$ | $34 \pm 14^\dagger$ | 0.03 |

<sup>†</sup>Extrapolated values

The product-dependent variation in apparent 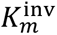 likely arose from different products reflecting a range of enzyme-surface interactions. The initial production rate of soluble aromatics reflects the generation of monomers and oligomers and subsequently reflects both complete and incomplete hydrolysis. On the other hand, LC-MS quantification of TPA reflects complete hydrolysis only. Consequently, the higher apparent 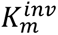 obtained from the fit of soluble aromatics reflects the fact that incomplete hydrolysis products (BHET, MHET and oligomers) result from less accessible surface sites that require greater enzyme loading to saturate. TPA-producing sites, being more accessible, saturate at lower enzyme concentrations and thus exhibit a lower apparent 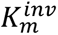. Despite this, both methods of analysis yielded apparent 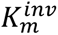 constants for CBD-PETase that were approximately 2-fold higher than those for HotPETase. The agreement between absorbance and LC-MS measurements supports the applicability of the inverse Michaelis-Menten model in capturing the kinetics of µPET depolymerization in this system. The two-fold higher apparent 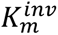 observed for CBD-PETase indicates that a greater enzyme concentration was required to achieve half-maximal enzymatic site occupancy compared to HotPETase. This strengthens the hypothesis that the CBM3a domain of the CBD-PETase enzyme engages in substrate competition with the HotPETase domain.^57–61^ The presence of two sites capable of binding µPET, only one of which can degrade µPET, increases the fraction of non-productive enzyme-µPET interactions. This provides a plausible explanation for why approximately twice the CBD-PETase concentration was required to reach comparable HotPETase product formation in free solution. This effect is expected to be reduced when CBD-PETase is immobilized on cellulose, as CBMs are known to preferentially bind to the cellulose substrate in the presence of PET,^62^ leaving the HotPETase domain as the only site available for µPET interactions.

On the other hand, both enzymes achieved comparable maximum reaction velocities. From the fit of TPA production, apparent 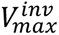 values of 1.45 ± 0.07 µM min⁻¹ and 2.05 ± 0.18 µM min⁻¹ were obtained for HotPETase and CBD-PETase, respectively. This was further supported by the fit of soluble aromatics production, which yielded apparent 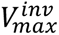 values of 1.27 ± 0.06 a.u. h^-1^ and 1.26 ± 0.05 a.u h^-1^ for HotPETase and CBD-PETase, respectively. These results indicate that both enzymes achieve similar enzymatic efficacies.^64^

Further kinetic assessment was performed by measuring the initial rate of product formation after incubation at 60 °C for 1 h as a function of increasing substrate concentration (0.75–25 g L⁻¹), with a fixed enzyme concentration of 0.45 µM. The conventional Michaelis-Menten model (SI Derivation 2) was then applied to quantitatively describe the system. Fitting the initial production rate of TPA yielded apparent 𝐾*_m_* values of 25 ± 11 g L⁻¹ and 34 ± 14 g L⁻¹, with corresponding apparent 𝑉*_max_* values of 16 ± 4 µM min⁻¹ and 16 ± 4 µM min⁻¹ for HotPETase and CBD-PETase, respectively. Fitting the initial production rate of soluble aromatics yielded apparent 𝐾*_m_* values of 22 ± 12 g L⁻¹ and 50 ± 40 g L⁻¹ with corresponding apparent 𝑉*_max_* values of 7 ± 2 a.u. h^-1^ and 8 ± 5 a.u. h^-1^ for HotPETase and CBD-PETase, respectively. Larger fitting errors were observed due to the inability to measure data points beyond the considerably high µPET concentration of 25 g L⁻¹, which falls below the predicted apparent 𝐾*_m_* of CBD-PETase. Despite this limitation, the conventional Michaelis-Menten kinetics also indicated that µPET depolymerization was limited by surface accessibility, as an excessively high substrate concentration (beyond practical limits) was required to saturate a modest enzyme concentration (0.45 µM). However, the similar 𝑉*_max_* obtained for both enzymes indicates similar enzymatic turnover for the two enzymes.

Finally, combining the inverse and conventional Michaelis-Menten parameters enabled estimation of the fraction of catalytically accessible surface sites, 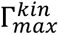, under the experimental conditions (SI Derivation 3). Both methods of characterizing product formation revealed a low fraction of accessible surface sites (∼2–4%, Table 1) in the high crystallinity µPET used in this study. There was not a discernible difference in accessibility for either enzyme, indicating that this limitation was driven by the µPET substrate, and that the carbohydrate binding domain likely did not interfere with access to the polymer surface. This quantifies an already established issue: current PETases struggle to depolymerize highly crystalline PET.^65,66^ As a result, research efforts have already begun to explore strategies to increase depolymerization of high crystallinity PET, including engineered PETases^35^ and pre-treatment^67^ of semi-crystalline PET. This strengthens the case for the development of an enzymatic filtration material, as not only could it incorporate improved PETases, but it could be treated during degradation to improve the depolymerization of high crystallinity µPET.

### Langmuir Binding Model Describes the Immobilization of CBD-PETase to Cellulose

Following characterization of the soluble enzymes, the ability of the CBD-PETase fusion to bind cellulose substrates was investigated. The CBM3a globular domain consists of a planar hydrophobic strip, composed of histidine, tryptophan, tyrosine and an arginine-aspartate ion pair on the lower binding surface.^42,43^ Surface mapping and mutagenesis studies have shown that the CBM3a binds to glucose molecules within a single chain of cellulose, as well as to those in adjacent cellulose chains to enhance the stability of the binding.^63^ Based on this binding mechanism, the immobilization of CBM3a to cellulose can be modeled by the Langmuir Adsorption model.^68–71^ Consequently, depletion assays were performed to determine the binding of CBD-PETase to cellulose at varying equilibrium concentrations, followed by fitting to the Langmuir model (SI Derivation 4). Three cellulose substrates, fibrous cellulose, microcrystalline cellulose and bacterial nanocellulose (BNC) from *K. xylinus*, were used to assess immobilization capacity.

Binding isotherms showed that CBD-PETase had the highest apparent adsorption capacity for bacterial nanocellulose followed by microcrystalline cellulose, then fibrous cellulose (Table 2). The order of magnitude higher adsorption capacity of BNC is hypothesized to be a result of both the high porosity and high crystallinity of the substrate. Control experiments using the HotPETase enzyme, lacking the cellulose binding domain, showed no significant binding to BNC from *K. xylinus* (SI Figure 5). This confirmed that the immobilization to BNC was due to the appended CBM3a domain, and that despite having a hydrophobic binding site for adhesion to PET, the HotPETase enzyme does not compete with the CBM3a domain for cellulose binding sites.

**Table 2.** Apparent *q*_max_ of CBD-PETase on Different Cellulose Substrates.

| Apparent $q_{\text{max}}$<br>CBD-PETase / cellulose (mg/mg) | | |
| --- | --- | --- |
| Fibrous cellulose | Microcrystalline cellulose | Bacterial nanocellulose |
| $0.013 \pm 0.001$ | $0.032 \pm 0.006$ | $0.2 \pm 0.06$ |

FT-IR was used to complement the binding analysis to confirm that protein was integrated in the BNC scaffold after immobilization. Analysis showed higher amide peak intensities (amide I: 1600–1700 cm^-1,^ amide II: ∼1550 cm^-1^), corresponding to the protein backbone, for the BNC-CBD-PETase composite compared to the BNC only control. FT-IR signal from the protein component could be detected at loading concentrations (*C*_i_) as low as 0.02 mg/mL (Figure 3c).

### BNC-CBD-PETase Composite Captures and Degrades µPET

Following demonstration that the CBD-PETase enzyme was catalytically active and could bind to cellulose, we next sought to demonstrate that the BNC-CBD-PETase composite could capture µPET during water filtration. Consequently, BNC-CBD-PETase composites were used to filter water containing 10 mg of µPET, followed by visual probing via scanning electron microscopy (SEM). SEM images revealed highly entangled nanofibers that formed pores of varying sizes (Scheme 1c). These pores were flexible and deformed around particles to enable the retention of irregularly shaped fragments and the structures were retained after post-growth cleaning, CBD-PETase immobilization, and µPET filtration (Figure 4a, SI Figure 6). Dynamic and hierarchical pore structures, such as those observed here, allow for the sequestration of microplastics of varying sizes, shapes, and structures.

**Figure 4.**
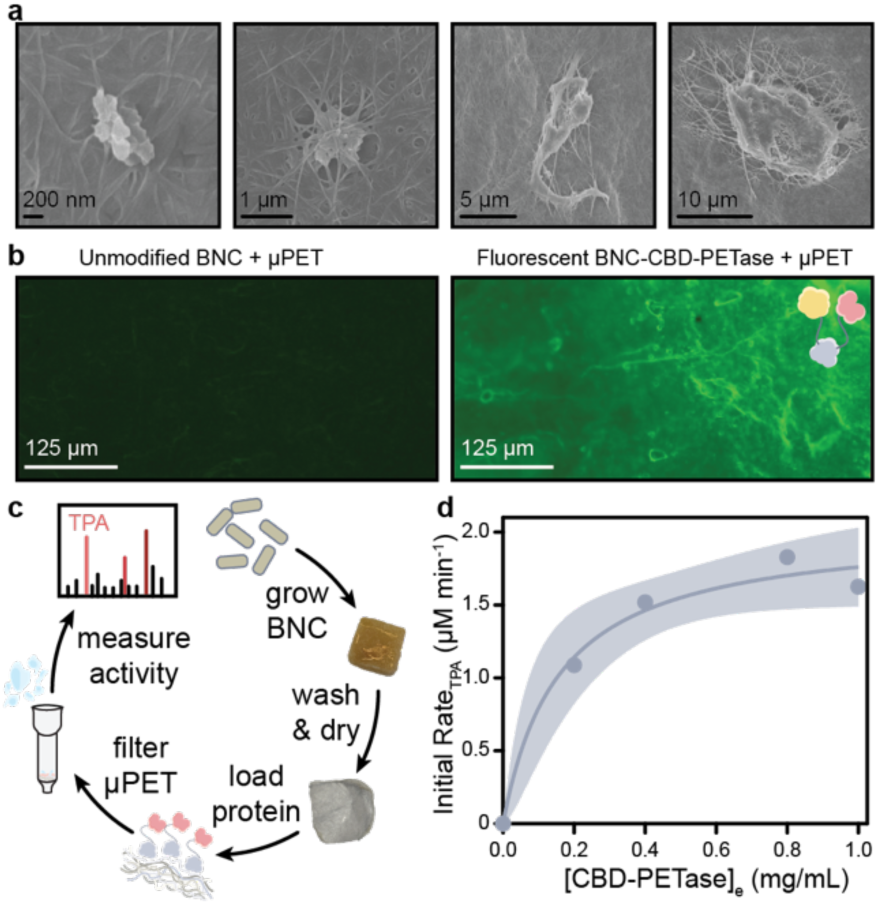
BNC-CBD-PETase Composites Capture and Degrade μPET. (a) SEM images showing μPET particles retained within the hierarchical nanofiber network of BNC-CBD-PETase composites. (b) Fluorescence microscopy images of unmodified BNC and fluorescently labeled BNC-CBD-PETase composite after μPET filtration (c) Schematic of the filtration and degradation process used to evaluate immobilized CBD-PETase activity. (d) Initial TPA production rate from BNC-CBD-PETase composites as a function of equilibrium CBD-PETase concentration after immobilization, 𝐶_*e*_, fitted to an inverse Michaelis-Menten analog. Data points indicate the average of 3 independent experiments, the line shows the fit to the appropriate model, and the shaded region corresponds to the 95% CI of the fit.

We then sought to demonstrate that the immobilized protein could interact with the trapped microplastics. Consequently, fluorescence microscopy, alongside a fluorescently labeled variant of the CBD-PETase enzyme (Venus-CBD-PETase), was used to visualize enzyme-plastic interactions. Venus-CBD-PETase was immobilized to purified BNC at a low enzyme loading (*C_i_* = 0.02 mg/mL) to prevent signal saturation, and then used to filter water containing 10 mg of µPET. An analogous control experiment, using purified BNC, was also conducted. Compared to the unmodified BNC control, the fluorescently labelled BNC-CBD-PETase composite exhibited a higher fluorescence intensity within the cellulose matrix. Additionally, even higher fluorescence was observed in areas where µPET was present, more pronounced than the intrinsic fluorescence of µPET plastics, as displayed in the unmodified BNC + µPET control (Figure 4b). These observations suggest that the enzyme was successfully retained within the scaffold after filtration and enriched to the microplastics captured within the composite.

Finally, to probe the catalytic activity of the immobilized enzyme, CBD-PETase was immobilized on a fixed amount of purified BNC (17 mm x 17 mm, 5 ± 0.5 mg), such that the BNC was at equilibrium with increasing concentrations of CBD-PETase. Consistent with the Langmuir binding model, this resulted in an increasing fraction of BNC sites immobilized with the CBD-PETase enzyme, and consequently, an increasing concentration of enzyme immobilized. Following enzyme immobilization, pellicles were used to filter water containing 10 mg of µPET, subjected to high temperature degradation for 1 h at 60 °C in Gly-OH (pH 9.2), followed by quantification of the initial TPA production rate via LC-MS analysis (Figure 4c). LC-MS quantification indicated that increasing concentrations of immobilized enzyme, as gauged by the *C*_e_ of the solution after immobilization, resulted in increased initial TPA production rate (Figure 4d).

Initial velocity results were fitted to an analog of the inverse Michaelis-Menten model to quantitatively describe the system. The apparent 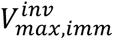 obtained from this model represents the maximum initial TPA production rate when 5 mg of BNC-CBD-PETase sequesters and degrades µPET under the experimental conditions. The apparent 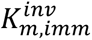 from this model represents the equilibrium CBD-PETase concentration, 𝐶*_e_*, required to produce a BNC-CBD-PETase composite with sufficient immobilized enzyme loading to reach half-maximal initial TPA production under the experimental conditions. Because 𝐶_*e*_ is related to 𝑞*_e_* through the Langmuir adsorption isotherm, it directly reflects the amount of CBD-PETase immobilized per mg of BNC-CBD-PETase composite.

From the fit, an apparent 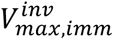 of 2.04 ± 0.18 µM min⁻¹ and an apparent 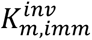 of 0.16 ± 0.06 mg/mL was obtained for 5 mg of BNC-CBD-PETase composite. In conjunction with the Langmuir analysis, this corresponded to ∼0.07 mg of immobilized enzyme and an effective enzyme concentration of 0.8 µM, giving a 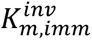 of the same order of magnitude as the free enzyme in solution 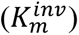.

### Symbiotic Co-Culture as a Route to Scalable, In Situ Production of Catalytically Active BNC-CBD-PETase Composites

We developed a co-culture strategy that provides a route towards scalable manufacture of the BNC-CBD-PETase composite. Yeast (*S. cerevisiae*) was engineered to secrete CBD-PETase during symbiotic growth with BNC-producing *K. xylinus*. This method enabled bottom-up assembly of the protein-cellulose composite, where the enzyme was immobilized during BNC biosynthesis (Figure 5a). FTIR analysis supports successful functionalization of BNC produced from the co-culture, with the composite material exhibiting more intense amide-associated features (N-H, C-N, and C=O) compared to the corresponding BNC control, produced from a co-culture consisting of the parent *S. cerevisiae* strain (Figure 5b). This indicates high CBM3a-mediated binding and retention during pellicle formation, despite the greater environmental complexity introduced by co-cultured growth.

**Figure 5.**
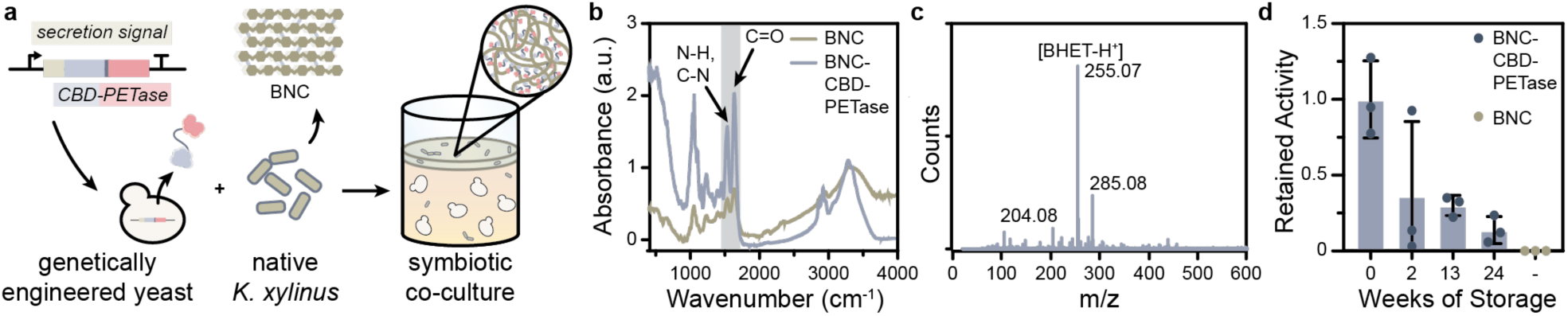
Symbiotic Co-Culture Enables In Situ Production of Catalytically Active BNC-CBD-PETase Composites. (a) Schematic depicting the use of a symbiotic co-culture, composed of genetically engineered yeast and *K. xylinus*, to produce the CBD-PETase BNC composite. (b) FT-IR analysis supports high CBD-PETase immobilization during cellulose biosynthesis due to the more dominant amide peaks from the immobilized protein. (c) Mass spectra highlighting the prominent BHET [M+H]⁺ peak observed after co-culture-derived composites were used to filter and degrade μPET. (d) Graph displays the fraction of activity maintained by the co-culture-derived composites after varying weeks of dry storage at ambient conditions. Individual data points are shown for n = 3; bars show the mean and error bars show the standard deviation.

Next, the catalytic functionality of the co-culture-derived BNC-CBD-PETase composite was evaluated. The co-culture-derived composite was used to filter water containing 10 mg of µPET, followed by degradation in Gly-OH (pH 9.2) for 6 h at 60 °C. An extended degradation time of 6 h was chosen to generate sufficient degradation products to dominate the background signal from residual media. LC-MS analysis of the smaller TPA fragment showed inconsistent observation of the characteristic [M+H]^+^ ion (m/z 167.03). This issue was again observed in control experiments, where known concentrations of TPA were analyzed in a matrix of Gly-OH and trace media from the co-culture. The similar inconsistencies observed in the controls suggest that detection of the TPA-associated ion was likely suppressed by residual components from the co-culture medium. In contrast, mass spectra consistently showed formation of BHET (Figure 5c), identified by the characteristic [M+H]^+^ ion (m/z 255.07). These results support that immobilized CBD-PETase remained catalytically active within the co-culture-derived composites.

The development of an active BNC-CBD-PETase composite through co-culturing was notable because it combines material synthesis and functionalization into a single biological process. In conventional immobilization approaches, enzymes are first produced and purified before being attached to a pre-formed material,^72^ requiring multiple processing and chemical modification steps that can increase production cost. Here, cellulose-producing *K. xylinus* and engineered *S. cerevisiae* work together to incorporate CBD-PETase directly into the growing BNC network during biosynthesis. Co-culture-derived BNC-CBD-PETase composites were then assessed for retained catalytic activity following dry storage at ambient conditions over 24 weeks. Activity was normalized to the BHET [M+H]⁺ ion peak area (m/z 255.07) obtained from freshly synthesized composites (0 weeks). Composites retained enzymatic activity for the full 24-week storage period (Figure 5d), demonstrating a level of robustness that supports their potential as shelf-stable, enzymatic filtration materials.

## Conclusion

In this work, we presented a biodegradable, catalytic material that combines the utility of bacterial nanocellulose and PETase, enabling both the capture and degradation of µPET in water. Structure-guided engineering was successfully used to modify a PETase variant for immobilization on bacterial nanocellulose. The effect of the additional domain on catalytic activity was quantitatively assessed by comparing it with the single-domain PETase enzyme, and we found that the additional cellulose-binding domain did not compromise activity. Instead, we found that the major limitation of the hybrid enzyme was low accessibility to surface sites within crystalline µPET, a limitation it inherited from the precursor HotPETase enzyme. Consequently, this system provides a foundation for the design of next-generation enzymatic materials, with possible improvements aligned with advances in PET hydrolases, as well as other plastic degrading enzymes, that overcome current limitations. Additionally, we successfully devised a co-culture strategy for in situ assembly of the composite catalytic material, providing a route for scalable material fabrication. Characterization of the catalytic activity of the co-culture-derived composites showed retained activity over extended storage periods at ambient conditions.

## Materials and Methods

### Media Preparation

For *E. coli* expression of recombinant enzymes, 2× YT was prepared using 16 g L⁻¹ casein hydrolysate (Thermo Scientific, AAJ12855P5), 10 g L⁻¹ yeast extract (Thermo-Fisher Scientific, J23547.A1), and 5 g L⁻¹ NaCl (Fisher Scientific, S271-3). Media were sterilized by autoclaving (30 min, liquid cycle), cooled, and supplemented with the appropriate antibiotic immediately prior to the addition of inoculum.

For bacterial nanocellulose production and co-culture growth, yeast peptone with dextrose (YPD) or sucrose (YPS) media were prepared with 10 g L⁻¹ yeast extract, 20 g L⁻¹ soy peptone (Millipore Sigma, 87972), and 20 g L⁻¹ dextrose (Sigma-Aldrich, G5767) or sucrose (Sigma-Aldrich, S1888), respectively, followed by autoclaving (30 min, liquid cycle).

For yeast selection, synthetic complete (SC) dropout media lacking uracil was prepared with 1.92 g L⁻¹ dropout supplement without uracil (Sigma-Aldrich, Y1501), 6.8 g L⁻¹ yeast nitrogen base (Sigma-Aldrich, Y0626), and 20 g L⁻¹ dextrose. Sugars and amino acids were autoclaved separately to prevent undesired Maillard reactions.

Solid media were prepared by adding 20 g L⁻¹ bacto-agar (BD, 214050) to the media prior to autoclaving when appropriate.

### Cloning

#### Genes

The gene sequences encoding Venus,^73^ CBM3a,^42,43^ and HotPETase^36^ were obtained from the literature, arranged as Venus-CBD-PETase (SI Table 1), and synthesized together as a single gene construct (Twist Bioscience) from which variants of the genes could be generated.

For yeast secretion, the gene sequence for the secretion signal for TFP19,^25,74^ alongside those used for screening (Mating Factor Alpha SP,^25^ Suc2 SP,^74^ Pir4,^75^ and MTLCC1 SP-Pro-peptide^25^), was obtained from the literature and synthesized with codon optimization for *S. cerevisiae* (Twist Bioscience). The empty vector backbone (pYTK001), Pre-Assembled URA3 Integration Vector (pYTK096), pTDH3 (pYTK009), Venus (pYTK033), and tTDH1 (pYTK056) were obtained from the MoClo-Yeast Toolkit (YTK) plasmid kit,^73^ a gift from John Dueber (Addgene kit # 1000000061).

#### Plasmid Construction

Primers were designed to amplify the gene of interest with ∼20 bp overlap. Primers (IDT) were resuspended in Milli-Q water and diluted to 10 μM. PCR reactions (50 μL total) contained: 10 ng template DNA, 1 μL forward primer, 1 μL reverse primer, 1 μL 10 mM dNTPs, 0.5 μL Phusion polymerase, 10 μL 5× HF buffer, and nuclease-free water. DNA amplification was performed with 35 cycles of denaturing (98 °C), annealing (primer-specific temperatures), and extension (72 °C). The annealing temperature and extension times were adjusted based on GC content and fragment length, respectively. DNA products were verified by agarose gel electrophoresis, and the fragments of interest were extracted and purified from the agarose gel with a QIAquick Gel Extraction kit, following the manufacturer’s instructions.

Plasmids for heterologous protein expression in *E. coli* were constructed by assembling purified PCR-amplified DNA fragments encoding the proteins of interest into purified PCR-amplified pET-21b(+) or pET28a(+) backbones using NEB HiFi Assembly, following the manufacturer’s instructions. HotPETase was cloned into pET-21b(+), whereas the fusion proteins CBD-PETase and Venus-CBD-PETase were cloned into pET28a(+). Resulting plasmids were transformed into *E. coli* NEB5α cells for plasmid amplification. After sequence verification (Genewiz), plasmids were transformed into NiCo21(DE3).

Plasmids for heterologous protein secretion in *S. cerevisiae* were generated using a combination of NEB HiFi Assembly and Golden Gate Assembly. To screen secretion capabilities under our experimental conditions, a library of MoClo-YTK-compatible Type 3a secretion signal parts was first generated by assembling genes encoding for secretion signal sequences (Mating Factor Alpha SP,^25^ Suc2 SP,^74^ TFP19,^25,74^ Pir4,^75^ and MTLCC1 SP-Pro-peptide^25^) into the pYTK001 backbone using NEB HiFi Assembly. Golden Gate Assembly was then used to combine these Type 3a secretion signal constructs with MoClo-YTK plasmid components pYTK096, pYTK009, pYTK045, and pYTK056, following the assembly protocol.^73^ After qualitative secretion screening, the plasmid construct containing the TFP19 secretion signal was selected for further studies.

The final secretion construct was generated using NEB HiFi Assembly. The selected plasmid was PCR-amplified, omitting the pYTK045 insert, to serve as the backbone, and the CBD-PETase gene was PCR amplified from the Venus-CBD-PETase fragment with backbone-compatible overhangs. Both fragments were gel-purified using a QIAquick Gel Extraction Kit. The fragments were then combined using NEB HiFi Assembly and transformed into *E. coli* NEB5α cells for plasmid amplification.

#### *E. coli* Transformation

Under sterile conditions, 2 μL of the plasmid-containing reaction mixture (from HiFi or Golden-Gate assembly) was added to a tube of thawed NEB 5-alpha competent *E. coli* cells. The mixture was gently flicked to mix, then incubated on ice for 30 min. Following incubation on ice, the mixture was heat-shocked in a 42 °C water bath for 30 s. Immediately after heat shock, the mixture was placed on ice for 5 min. The cells were then recovered by adding 950 μL of pre-warmed SOC media (NEB, B9035) and then placed in a 37 °C incubator with shaking for 1 h. After the incubation, the cells were plated on solid LB media containing the appropriate selection antibiotic. Individual colonies from the plates were grown in liquid LB media containing the appropriate antibiotic for 16-18 h. A portion of the culture was stored as a 40% glycerol stock at -80 °C, and the remaining portion was used to isolate plasmids (Qiagen, QIAprep spin miniprep kit). Isolated plasmids were sequence verified by whole plasmid sequencing (Genewiz).

Sequence-verified plasmids assembled for expression in *E. coli* were transformed into competent NiCo21(DE3) cells using the same protocol.

#### S. cerevisiae Transformation

Plasmids for yeast chromosomal integration were linearized by digestion with NotI-HF (NEB, R3189S) using the manufacturer’s protocol, followed by transformation into *S. cerevisiae* strain BY4733 (MATa his3Δ200 leu2Δ0 met15Δ0 trp1Δ63 ura3Δ0).

Yeast Transformation Kit (Sigma-Aldrich, YEAST1-1KT) was purchased to supply reagents for yeast transformation. *S. cerevisiae* strain BY4733 (MATa his3Δ200 leu2Δ0 met15Δ0 trp1Δ63 ura3Δ0) was plated on YPD agar plates and grown for 4 d at 30°C or until colonies appeared. A single colony was used to inoculate 5 mL of YPD and grown for 24 h at 30 °C with shaking at 250 rpm. The culture was then diluted to OD600 ∼0.3 and re-grown to mid-log phase (OD600 < 0.7). Cells were then harvested by centrifugation at 4000 rpm for 5 min and resuspended in 0.5 mL of transformation buffer (Sigma-Aldrich, T0809). Resuspended cells were centrifuged for 1 min using a micro-centrifuge, after which 0.45 mL of the supernatant was removed from the tube. To the tube, 10 μL of 10 mg/mL salmon testes DNA (Sigma-Aldrich, D9156) was added, followed by 1 µg of linearized plasmid. The mixture was then mixed by vortexing for 10 s. Next, 600 μL of Plate Buffer (Sigma-Aldrich, P8966) was added, followed by a second round of mixing by vortexing for 10 s. The mixture was incubated for 30 min at 30 °C with shaking at 250 rpm, then DMSO was added (NEB, B0515) to 10% (v/v). The mixture was subjected to an extended 15 min heat shock at 42 °C, followed by a spin-down using a mini-centrifuge for 3 s to facilitate removal of the supernatant. The pellet was then resuspended in 200 µL of Milli-Q water and then transferred to plates of SC dropout media lacking uracil. Plates were incubated at 30 °C until colonies formed (∼3–5 d). Chromosomal integration was confirmed by colony PCR.

### Protein Expression and Purification

#### Expression

All 2× YT media used were supplemented with 100 µg of the appropriate antibiotic (Ampicillin or Kanamycin) based on the expression plasmid. A NiCo21(DE3) *E. coli* colony containing the expression plasmid was used to inoculate 5 mL of 2× YT, followed by overnight growth (16-18 h) at 37 °C with shaking at 200 rpm. The resulting culture was then used to inoculate a 2.8 L Fernbach flask containing 1 L of 2× YT. Cultures were grown to OD600 ∼ 0.8, induced with 1 mL of 1 M Isopropyl β-D-1-thiogalactopyranoside (IPTG), and subsequently grown at 27 °C for ∼15 h post-induction. Cells were harvested by centrifugation at 4000 rpm for 10 min and stored at -80 °C until purification.

#### Purification

Cell pellets from 1 L cultures were resuspended in 30 mL lysis buffer (30 mM imidazole, 50 mM sodium phosphate, 300 mM NaCl, pH 8), followed by sonication on ice with a 0.5 in probe at 40% amplitude, with pulses of 2 s on/4 s off for 10 min. Lysate was clarified by two rounds of centrifugation at 4 °C and 10,000 rpm for 30 min and applied to a Ni-NTA column. Protein was purified using an ÄKTA system, where it was washed with 5 column volumes (CVs) of lysis buffer and eluted with elution buffer (250 mM imidazole, 50 mM sodium phosphate, 300 mM NaCl, pH 8). Proteins were stored in elution buffer at 4 °C and dialyzed into 50 mM Gly-OH buffer, pH 9.2 prior to use.

### Bacterial Nanocellulose (BNC) Pellicle Production

The cellulose-producing strain*, K. xylinus,* was first plated on YPD-agar and grown for 5–7 d at 30 °C or until individual colonies appeared. A single colony was then used to inoculate 5 mL of YPD media supplemented with 50 µL of cellulase (Sigma-Aldrich, C2730) and then grown at 30 °C for 5 d. After growth, the OD600 was measured to determine the appropriate dilution factor. Cell pellets were harvested by centrifugation at 3200 ×g for 10 min and resuspended in fresh YPD media (for BNC pellicles used for Ex Situ Enzyme Immobilization) or YPS (for <u>BNC pellicles</u> <u>derived from Co-Culture)</u> to an OD600 = 2.5 to remove cellulase.

#### BNC Pellicles used for Ex Situ Enzyme Immobilization

BNC pellicles were prepared by combining 200 μL of resuspended *K. xylinus* in YPD at an OD600 = 2.5 and 5 mL of cellulase-free YPD in 24-deepwell plates (Corning, P-DW-10ML-24-C-S). Cultures were grown statically for 5 d at 30 °C to allow BNC pellicle formation.

BNC pellicles were harvested and treated to remove trace media and cell particulate. This was accomplished through three rounds of washing with 0.1 N NaOH at 60 °C, followed by washing with excess Milli-Q water under vacuum using a Büchner funnel. Pellicles were dried at ambient conditions, then weighed to determine cellulosic mass.

#### BNC Pellicles derived from Co-culture

The CBD-PETase-producing *S. cerevisiae* strain, alongside an un-engineered counterpart (BY4733), was first plated on YPD-agar and grown for 5 d at 30 °C. A single colony was then used to inoculate 5 mL of YPD media. Cultures were grown for 18 h at 30 °C with shaking at 250 rpm and diluted to OD600 ∼ 0.01 in YPS media.

Co-cultures were produced by combining 200 μL of resuspended *K. xylinus* in YPS at an OD600 = 2.5, 200 μL of the appropriate *S. cerevisiae* strain in YPS at an OD600 ∼ 0.01, and 5 mL YPS in 24-deepwell plates. To obtain a co-culture-produced BNC pellicle devoid of CBD-PETase the parent BY4733 yeast strain was used. Cultures were grown statically for 5 d at 30 °C to allow BNC pellicle formation.

BNC pellicles were harvested and washed with excess Milli-Q water under vacuum using a Büchner funnel, to remove as much residual media and particulate matter as possible. A NaOH wash was not utilized to avoid disruption of the immobilized protein. BNC pellicles were dried under ambient conditions and stored at room temperature until use.

### Ex Situ Protein Immobilization on cellulose

Incubation utilized a known initial protein concentration (*C*_i_), as determined by Quant-iT™ Protein Assay Kit in conjunction with a sigmoidal four-parameter logistic (4PL) fit (GraphPad Prism).

Preliminary experiments showed minimal changes in supernatant concentration after 24 h of protein-cellulose incubation at room temperature; therefore, we assumed that *C*_e_ is equivalent to the supernatant concentration after 24 h of incubation with 1 mL of protein of initial concentration *C*_i_ and a fixed amount of cellulose. Post incubation, samples were centrifuged for 10 min at 10,000s×g using a tabletop centrifuge, and the protein concentration of the supernatant was determined by Quant-iT™ Protein Assay Kit using a sigmoidal 4PL curve. Three independent samples were prepared, and the average was used to determine the *C*_e._ Depletion of protein (*C*_i_ − *C*_e_), normalized to the mass of cellulose, was used to determine *q*_e_ (mg of protein bound / mg of cellulose) at each *C*_e_ concentration.

Exactly 6.5 mg of Fibrous Cellulose (Sigma Aldrich, C6288) or Microcrystalline cellulose (Sigma Aldrich, 435246) were used for each experiment, and each experiment was done in triplicate. BNC pellicles, on the other hand, had varying weights (between 4.5-5.5 mg) due to small variations from *K. xylinus* production; however, BNC pellicles maintained a constant dimension of 17 mm x 17 mm (L x W) as dictated by the shape of the wells in which they were produced.

### Assessing µPET Degradation of Free Enzymes

All free enzyme µPET degradation experiments were conducted in 1 mL volumes and in triplicate. Each experiment utilized 0.45 µM of enzyme and 2 g L⁻¹ μPET (50 mM Gly-OH, pH 9.2), with the exception of the inverse and conventional Michaelis-Menten experiments, in which enzyme and µPET concentrations were varied, respectively.

Preliminary characterization experiments probing temperature-dependent product formation were incubated at 20 °C, 40 °C, and 60 °C and analyzed after 3 h. Preliminary experiments probing time-dependent product formation were continuously monitored over a 5 h incubation at 60 °C. All inverse and conventional Michaelis-Menten experiments were incubated at 60 °C and analyzed after 1 h (during the linear initial velocity regime).

All experiments were analyzed using UV-Vis spectroscopy (Agilent, Cary60). Samples from experiments requiring the quantification of TPA produced were centrifuged at 10,000 ×g for 30 min at 25 °C after incubation to obtain the supernatant, which was then analyzed by LC-MS using an Agilent 1260 HPLC and 6230 TOF with the Agilent MassHunter Quantitative Analysis software (Version 11.0).

### Assessing µPET Degradation of Immobilized Enzymes

BNC-CBD-PETase composite pellicles were placed at the base of a Bio-Rad poly-prep column. To the pellicle, 5 mL of µPET (2 g L⁻¹) in water was added and filtered under gravity. BNC pellicles were removed from the column, followed by incubation in 2 mL of 50 mM Gly-OH, pH 9.2, at 60 °C.

#### BNC-CBD-PETase Pellicles from Ex Situ Immobilization

BNC-CBD-PETase pellicles were obtained using the **Ex Situ Protein Immobilization** method, as described earlier, and the *C*_e_ for each pellicle was determined. BNC-CBD-PETase pellicles were placed at the base of a Bio-Rad poly-prep column and used to gravity-filter 5 mL of µPET (2 g L⁻¹) in Milli-Q water.

Following filtration, BNC-CBD-PETase pellicles were incubated in 2 mL of Gly-OH (50 mM, pH 9.2), for 1 h (initial velocity regime of the free enzyme). The concentration of TPA generated per min was determined by LC-MS analysis.

The data were fitted to an analog of the inverse Michaelis-Menten model, where the y axis displays the initial rate of TPA generated and the x axis displays *C*_e._ The y-axis reflects the initial concentration of TPA generated per min when a fixed amount of BNC-CBD-PETase (L x W: 17 mm x 17 mm, mass: ∼5 ± 0.5 mg) was used to sequester microplastics from 5 mL of µPET (2 g L⁻¹) in Milli-Q water, and subjected to incubation at 60 °C in 2 mL of Gly-OH (50 mM, pH 9.2).

The x-axis reflects the concentration of the free enzyme that must be at equilibrium with any mass of BNC after immobilization, to obtain a fixed fractional occupancy of the CBD-PETase ligand on BNC binding sites. Since experiments were conducted with effectively a fixed amount of BNC-CBD-PETase pellicle (17 mm x 17 mm, mass: ∼5 ± 0.5 mg), *C*_e_ is directly related to the concentration of CBD-PETase enzymes present in each BNC pellicle, and thus effectively represents the enzyme concentration as required for the inverse Michaelis-Menten model.

The 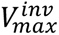 obtained from this model therefore represents the maximum initial concentration of TPA generated per minute when a fixed amount of BNC-CBD-PETase (L x W: 17 mm x 17 mm, mass: ∼5 ± 0.5 mg) was used to sequester microplastics from 5 mL of µPET (2 g L⁻¹) in Milli-Q water, and subjected to incubation at 60 °C in 2 mL of Gly-OH (50 mM, pH 9.2). The 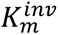 on the other hand, represents the *C*_e_ needed to produce a 5 mg BNC-CBD-PETase composite where half-maximal initial TPA production is achieved when used to sequester microplastics from 5 mL of µPET (2 g L⁻¹) in Milli-Q water, and subjected to incubation at 60 °C in 2 mL of Gly-OH (50 mM, pH 9.2).

#### BNC-CBD-PETase Composite Pellicles from In Situ Immobilization (Co-Culture)

BNC-CBD-PETase pellicles were created in situ during co-culturing with secreted enzyme immobilized during biosynthesis. After production, the pellicles were harvested, washed, dried and stored at room temperature until use, as described above.

BNC-CBD-PETase pellicles were placed at the base of a Bio-Rad poly-prep column and used to gravity-filter 5 mL of µPET (2 g L⁻¹) in Milli-Q water. Pellicles were then incubated at 60 °C in 2 mL of Gly-OH buffer (50 mM, pH 9.2) for 6 h. Extended incubation was performed to generate sufficient product to overcome the background LC-MS signal caused by residual complex media from the co-culture. Experiments were conducted in triplicate.

Samples containing cellulose were not analyzed by UV-Vis due to the inherent absorbance of cellulose at the wavelength used to analyze PET degradation products (λ = 242 nm).

### LC-MS Analysis

Samples were centrifuged (10,000 ×g, 30 min) to remove particulate matter prior to injection. LC-MS analysis was performed using a ZORBAX-Extend *C18* (1.8 µm, 2.1 x 50 mm) column with a mobile phase consisting of water (0.1% formic acid) and acetonitrile (0.1% formic acid).

TPA Calibration standards were generated by pre-dissolving Terephthalic acid (Sigma-Aldrich, 185361) in 100 mM phosphate buffer (K_2_HPO_4_-KH_2_PO_4,_ pH 9), to aid dissolution, followed by dilution in Milli-Q water to a final concentration of 10 mM TPA in 1 mM phosphate. The 10 mM TPA stock was further diluted to 1 mM in Gly-OH (50 mM, pH 9.2). The 1 mM stock solutions, alongside Gly-OH (50 mM, pH 9.2) was then used to prepare calibration standards in the range 5 µM-80 µM such that the solution was >99% Gly-OH and the final phosphate concentration was < 8 µM.

BHET Calibration standards were generated by dissolving 10 mM Bis(2-hydroxyethyl) terephthalate (Sigma-Aldrich, 465151) in Milli-Q water to aid dissolution, followed by further dilution to 1 mM in Gly-OH (50 mM, pH 9.2). The 1 mM stock solution, alongside Gly-OH (50 mM, pH 9.2) was then used to prepare calibration standards in the range of 5 µM - 80 µM such that the solution was >99% Gly-OH.

An LC-MS acquisition method was developed to elute TPA and BHET calibration standards at different retention times, such that the signal of the monomer’s characteristic [M+H]^+^ m/z ion that appeared at said retention time was linear with respect to concentration.

### FT-IR Analysis

Composite pellicles and corresponding BNC control pellicles produced by similar methods and subjected to the same cleansing steps were dried and their composition observed by FTIR (FTIR-ATR, Spectrum 100, Perkin Elmer, Waltham, MA). Absorbance spectra were collected with 100 scans with a spectral resolution of 4 cm^-1^. Samples were normalized to the characteristic hydroxyl peak of cellulose (3650-3350 cm^-1^) and the amide peak intensities (amide I: 1600-1700 cm^-1^ and amide II: ∼1550 cm^-1^) compared to the corresponding BNC pellicle controls, devoid of CBD-PETase immobilization.

### SEM

Representative samples of BNC pellicles functionalized with CBD-PETase before and after filtration of a 2 g L^-1^ µPET solution were dried at room temperature. The samples were attached to metal stubs using copper tape, sputter-coated with 4 nm of 5% Pd/Au (Cressington 108 Auto Sputter Coater) and imaged using a Zeiss Sigma VP Scanning Electron Microscope.

### Fluorescence Microscopy

Representative samples of BNC pellicles treated with a solution of Venus-CBD-PETase (*C*_i_ = 0.02 mg/mL), and an unfunctionalized BNC pellicle control were subjected to filtration and dried at room temperature. Pellicles were placed on microscope slides, followed by a drop of Milli-Q water and a coverslip to secure the sample. Microscopic images were obtained using an EVOS™ FL Auto 2 Imaging System using a 20× 0.4 NA Fluor objective, with both brightfield and GFP (Excitation: 470/22 nm, Emission: 525/50 nm) channels.

### DSC

A known mass of the µPET used in this study was placed in a Tzero pan and sealed with a lid using a Tzero press. The mass was recorded in the TRIOS software, and the sample analyzed using a TA Instruments Discovery DSC 250. The crystallinity and T_g_ were determined using the TRIOS software.

## Author Contributions

**Brittany Pitt:** Conceptualization (equal); Methodology, Investigation, Formal Analysis, Data Curation, Visualization, Project Administration (lead); Writing-Original Draft (lead); Writing-Review & Editing (equal). **Allie Obermeyer**: Supervision (lead); Funding Acquisition (lead); Resources (lead); Conceptualization (equal); Writing-Reviewing & Editing (equal). **Dayana Guillen**: Investigation, Writing-Review & Editing (supporting).

## Notes

The authors declare no competing financial interest.

## Acknowledgements

We gratefully acknowledge Alana Kwan and Leslie Ramirez for help with data collection and lab experiments, Natalie Ling for cryo-milling µPET from Goodfellow PET granules, Zubin Kumar for DSC characterization of cryo-milled µPET, and Thomas Bina for support with FT-IR measurements.

