## Supplementary Information for "Catalytic Bacterial Nanocellulose Composite That Captures and Degrades PET Microplastics"

### Supporting Information

**Towards Closing the Loop on Microplastic Filtration: A Catalytic Bacterial Nanocellulose Composite That Captures and Degrades PET Microplastics**

**Brittany N. Pitt,[1] Dayana Guillen,[1] and Allie C. Obermeyer[1]***

1. Department of Chemical Engineering, Columbia University, New York, NY 10027

*

| Protein | Amino Acid Sequence | MW (Da) |
|---|---|---|
| HotPETase[1] | MQTNPYARGPNPTAASLEASAGPFTVRSFTVARPVGYGAGTVYYP<br>TNAGGTVGAIAIVPGYTATQSSINWWGPRLASHGFVVITIDTNSTLD<br>KPESRSSQQMAALRQVASLNGTSSSPIYGKVDTARGGVMGWSMG<br>GGGSLISAANNPSLKAAAVMAPWHSSTNFSSVTVPTLIFACENDRI<br>APVKEYALPIYDSMSLNAKQFLEICGGSHSCACSGNSNQALIGMKG<br>VAWMKRFMDNDTRYSQFACENPNSTAVCDFRTANCSLEHHHHHH | **28,774.6** |
| CBD[2,3]-PETase | MGHHHHHHGGANATPTKGATPTNTATPTKSATATPTRPSVPTNTPT<br>NTPANTPVS**GNLKVEFYNSNPSDTTNSINPQFKVTNTGSSAIDLS**<br>**KLTLRYYYTVDGQKDQTFWCDHAAIIGSNGSYNGITSNVKGTF**<br>**VKMSSSTNNADTYLEISFTGGTLEPGAHVQIQGRFAKNDWSNY**<br>**TQSNDYSFKSASQFVEWDQVTAYLNGVLVWGKEP**GGSVVPSTQ<br>PVTTPPATTKPPATTKPPATTIPPSDDPGGGGGG**MQTNPYARGPNPT**<br>**AASLEASAGPFTVRSFTVARPVGYGAGTVYYPTNAGGTVGAIAI**<br>**VPGYTATQSSINWWGPRLASHGFVVITIDTNSTLDKPESRSSQQ**<br>**MAALRQVASLNGTSSSPIYGKVDTARGGVMGWSMGGGGSLIS**<br>**AANNPSLKAAAVMAPWHSSTNFSSVTVPTLIFACENDRIAPVKE**<br>**YALPIYDSMSLNAKQFLEICGGSHSCACSGNSNQALIGMKGVA**<br>**WMKRFMDNDTRYSQFACENPNSTAVCDFRTANCS** | **54,412.1** |
| Venus[4]-CBD-PETase | MGHHHHHHGGA**SKGEELFTGVVPILVELDGDVNGHKFSVSGEG**<br>**EGDATYGKLTLKLICTTGKLPVPWPTLVTTLGYGLQCFARYPD**<br>**HMKQHDFFKSAMPEGYVQERTIFFKDDGNYKTRAEVKFEGDT**<br>**LVNRIELKGIDFKEGGNILGHKLEYNYNSHNVYITADKQKNGIK**<br>**ANFKIRHNIEDGGVQLADHYQQNTPIGDGPVLLPDNHYLSYQS**<br>**ALSKDPNEKRDHMVLLEFVTAAGITHGMDELYK**NATPTKGATP<br>TNTATPTKSATATPTRPSVPTNTPTNTPANTPVS**GNLKVEFYNSNPS**<br>**DTTNSINPQFKVTNTGSSAIDLSKLTLRYYYTVDGQKDQTFWC**<br>**DHAAIIGSNGSYNGITSNVKGTFVKMSSSTNNADTYLEISFTGG**<br>**TLEPGAHVQIQGRFAKNDWSNYTQSNDYSFKSASQFVEWDQV**<br>**TAYLNGVLVWGKEP**GGSVVPSTQPVTTPPATTKPPATTKPPATTIPP<br>SDDPGGGGGG**MQTNPYARGPNPTAASLEASAGPFTVRSFTVARP**<br>**VGYGAGTVYYPTNAGGTVGAIAIVPGYTATQSSINWWGPRLAS**<br>**HGFVVITIDTNSTLDKPESRSSQQMAALRQVASLNGTSSSPIYGK**<br>**VDTARGGVMGWSMGGGGSLISAANNPSLKAAAVMAPWHSST**<br>**NFSSVTVPTLIFACENDRIAPVKEYALPIYDSMSLNAKQFLEICG**<br>**GSHSCACSGNSNQALIGMKGVAWMKRFMDNDTRYSQFACENP**<br>**NSTAVCDFRTANCS** | **80,965.4** |

**SI Table 1. Amino acid sequences of HotPETase variants.** Domain names and sequences are highlighted using the same color. All globular domains are written in bolded letters, whereas linkers and tags are written in plain script. Purification tags with linkers and cut sites are highlighted in yellow, and the added flexible linker is highlighted dark grey. The native CBM3a linkers are highlighted with the same color as the CBD domain but is written in plain script.

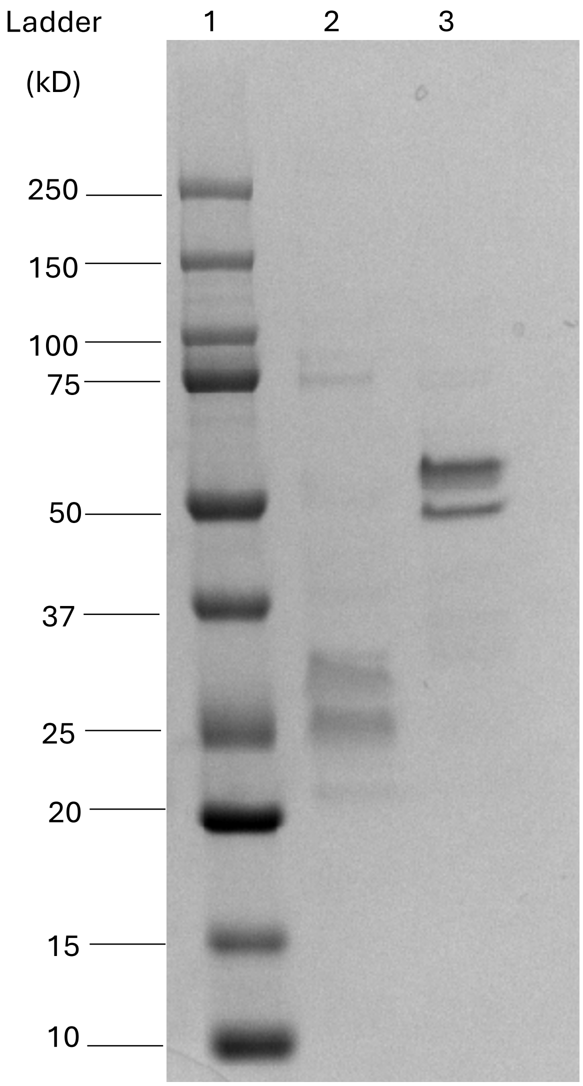

**SI Figure 1. SDS PAGE of PETase variants.** Purified enzymes used for quantitative studies were analyzed by SDS-PAGE to confirm molecular weight (MW) and purity. Lane 1: ladder, lane 2: HotPETase (expected MW ~28.7 kDa), and lane 3: CBD-PETase (expected MW ~ 54.4kDa).

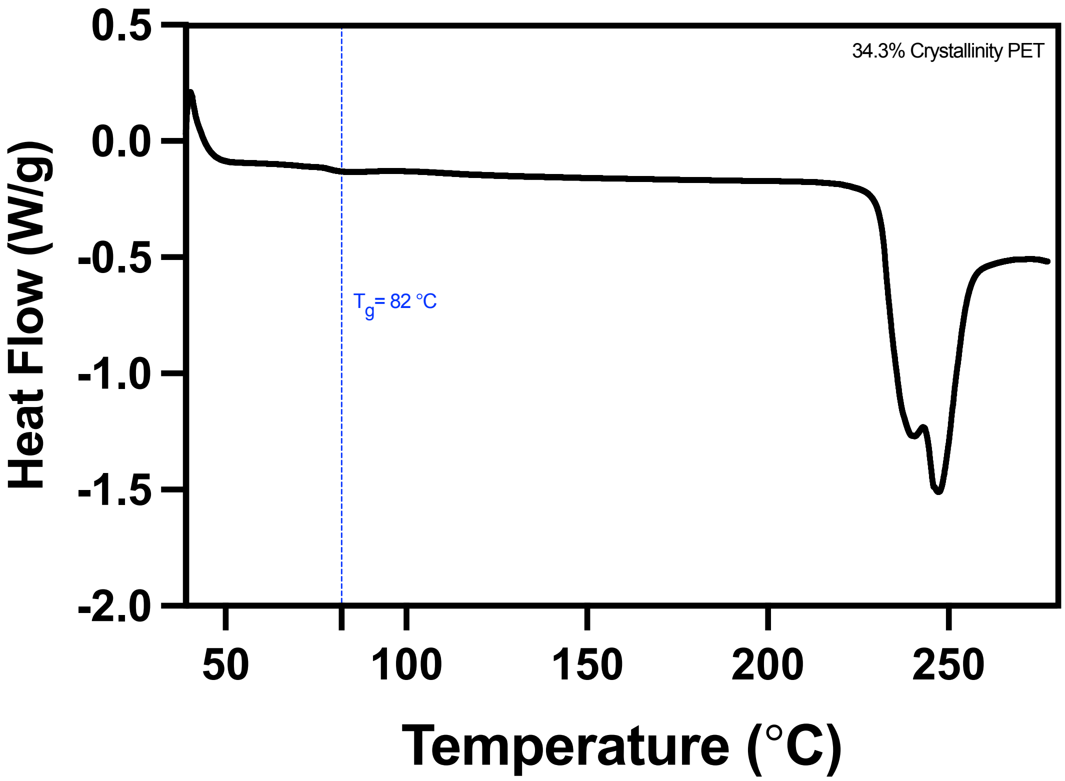

**SI Figure 2. DSC of μPET.** DSC scan of the μPET used in this study confirms crystallinity is retained after cryo-milling PET granules to obtain microplastics. The $T_g$ of the μPET was 82 °C, with a crystallinity of 34.3%[5].

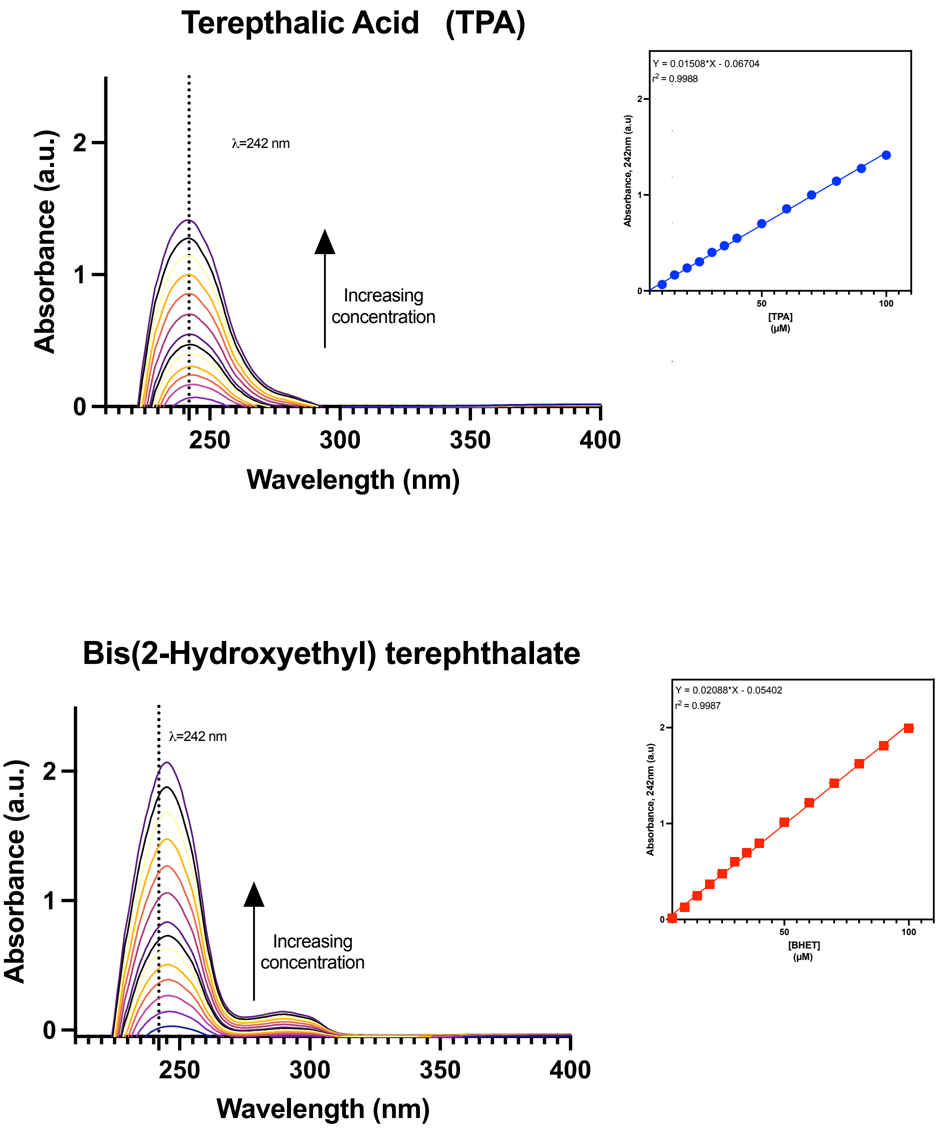

**SI Figure 3. UV-Vis calibration of BHET and TPA.** The absorbance from 200-400 nm of varying concentrations (0-100 μM) of TPA and BHET in 50 mM GlyOH buffer, pH 9.2 was measured. Linear regression (GraphPad Prism 11) of the absorbance at 242 nm as a function of TPA or BHET concentration demonstrates a clear linear correlation between absorbance and concentration[6,7] for both PET degradation products assessed in this study.

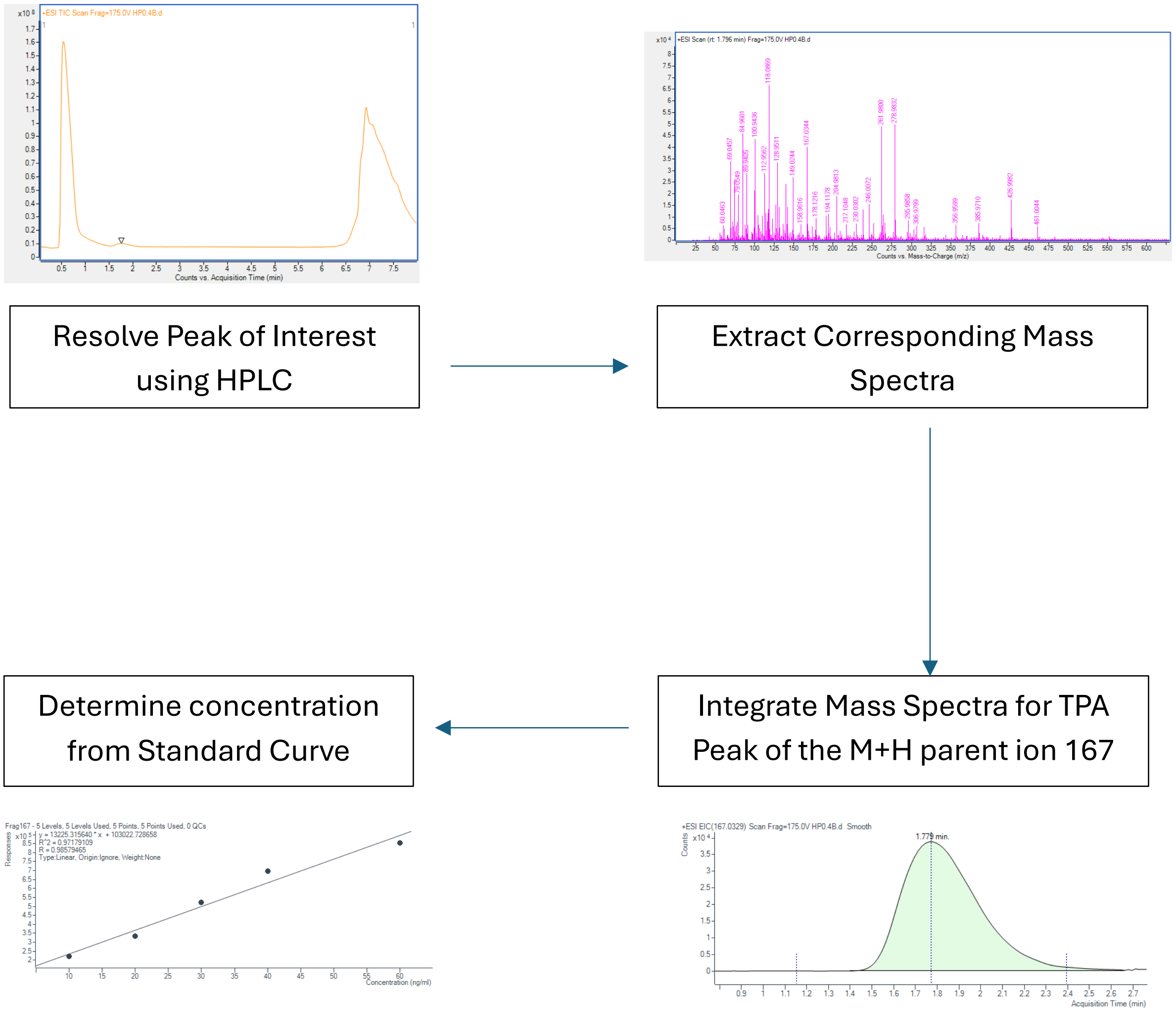

**SI Figure 4. Example workflow of LC/MS-based monomer concentration determination.** The liquid chromatography method[7,8] was optimized to elute TPA at a retention time of 1.8 min. The corresponding mass spectra at this time were then extracted, from which the $[M+H]^+$ peak[9] (167.03 m/z) was integrated. The integration of this peak was then compared to that of the standard TPA curve, which shows a linear response to increasing TPA concentration, with an $r^2$=0.97.

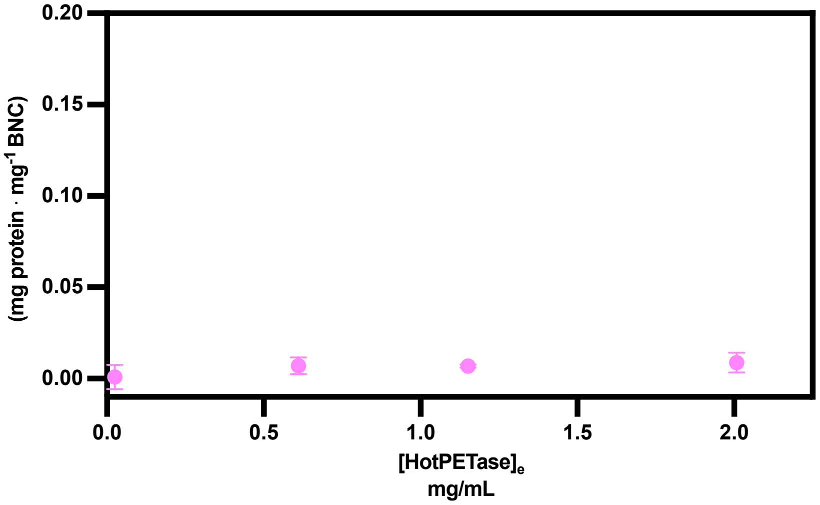

**SI Figure 5. Immobilization of HotPETase on Bacterial Nanocellulose (BNC).** Depletion assay confirms that the catalytic HotPETase domain does not significantly bind to Bacterial Nanocellulose (BNC) in the absence of a cellulose-binding domain (CBD).[10,11]

Pellicle Processing for Immobilization

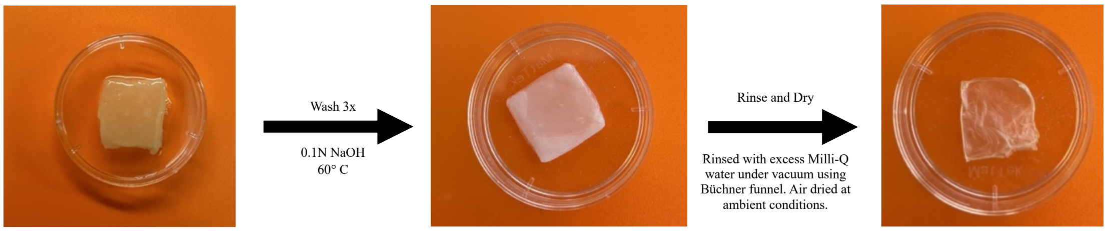

Pellicle Processing for Co-culture grown Pellicles

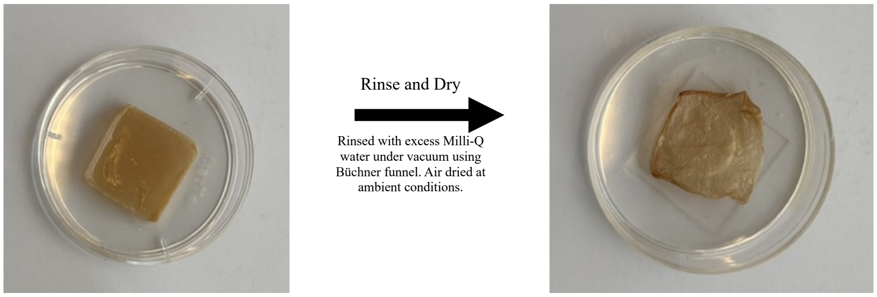

**SI Figure 6. Schematic of BNC Pellicle processing for downstream applications**. Pellicles used for immobilizing proteins are completely stripped from possible interference using multiple washes of 0.1 N NaOH at 60 °C, followed by a rinse of approximately 250 mL per 17mmx17mm pellicle. Pellicles from co-culture were solely rinsed with excess water to prevent interference with the in situ immobilized proteins.

$$E + S^* \underset{k_{-1}}{\overset{k_1}{\rightleftharpoons}} ES^* \xrightarrow{k_{cat}} E + P$$

$S^*$ :accessible surface sites

**Assumptions:**

1. Initial rate conditions ($[P] \approx 0$
2. Steady-state: $\frac{d[ES*]}{dt} \approx 0$
3. Accessible Substrate conservation: $[S^*]_T = [S^*] + [ES^*]$ **(1)**

**Steady-state for $ES^*$**

$$0 = k_1[E][S^*] - (k_{-1} + k_{cat})[ES^*]$$

Define $K_M^{inv} \equiv \frac{k_{-1}+k_{cat}}{k_1}$,

$$[ES^*] = \frac{[E][S^*]}{K_M^{inv}} \quad \textbf{(2)}$$

**Substitute equation (1) into equation (2)**

$[S^*] = [S^*]_T - [ES^*]$

$$[ES^*] = \frac{[E]([S^*]_T - [ES^*])}{K_M^{inv}}$$

$$[ES^*] = \frac{[S^*]_T[E]}{K_M^{inv}+[E]} \quad (3)$$

**Initial Velocity**

$$v = k_{cat}[ES^*]$$

$$v = \frac{k_{cat}[S^*]_T\,[E]}{K_M^{inv}+[E]} \quad (4)$$

**Define $V_{max}^{inv}$**

At high [E] all accessible sites are occupied: $[ES^*] \rightarrow [S^*]_T$

$$V_{max}^{inv} \equiv k_{cat}[S^*]_T \quad \textbf{(5)}$$

$$\boxed{v = \frac{V_{max}^{inv}\,[E]}{K_M^{inv}+[E]}} \quad (6)$$

**SI Derivation 1. Inverse Michaelis-Menten equation.**[12]

$$E + S \underset{k_{-1}}{\overset{k_1}{\rightleftharpoons}} ES \xrightarrow{k_{cat}} E + P$$

Assumptions

1. Initial rate conditions ($[P] \approx 0$
2. Steady-state: $\frac{d[ES]}{dt} \approx 0$
3. Enzyme conservation: $[E]_T = [E] + [ES]$ **(7)**

**Steady-state for *ES***

$$0 = k_1[E][S] - (k_{-1} + k_{cat})[ES]$$

Define $K_M = \frac{k_{-1}+k_{cat}}{k_1}$,

$$[ES] = \frac{[E][S]}{K_M} \quad \textbf{(8)}$$

**Substitute equation (6) into equation (7)**

$[E] = [E]_T - [ES]$

$$[ES] = \frac{[E]_T[S]}{K_M+[S]} \quad \textbf{(9)}$$

**Initial velocity**

$v = k_{cat}[ES]$

$$v = \frac{k_{cat}[E]_T[S]}{K_M+[S]} \quad \textbf{(10)}$$

**Define $V_{max}$**

$$V_{max} = k_{cat}[E]_T \quad \textbf{(11)}$$

$$\boxed{v = \frac{V_{max}[S]}{K_M+[S]}} \quad \textbf{(12)}$$

**SI Derivation 2. Michaelis-Menten equation.**[13–17]

**Conventional Michaelis–Menten**

$$V_{max} = k_{cat}[E]_T \quad \textbf{(11)}$$

**Inverse Michaelis–Menten**

$$\mathrm{V_{max}^{inv}} = \mathrm{k_{cat}}[\mathrm{S}^*]_\mathrm{T} \quad \textbf{(5)}$$

Since Inverse MM does not assume that the substrate mass,

which is held constant, equals the total number of accessible attack sites,

$$[\mathrm{S}^*]_\mathrm{T} = \Gamma_{\mathrm{max}}^{kin} S_m$$

Where:

$[\mathrm{S}^*]_\mathrm{T}$ = total accessible attack sites

$S_m$= substrate mass concentration

$\Gamma_{\mathrm{max}}^{kin}$ = fraction of sites available for catalysis

Therefore, the equation can be rewritten as

$$\mathrm{V_{max}^{inv}} = \mathrm{k_{cat}}[\Gamma_{\mathrm{max}}^{\mathrm{kin}} S_m ] \quad \textbf{(5b)}$$

**Fraction of Substrate Available for Catalysis**

Divide equation (5b) by equation (11)

$$\Gamma_{\mathrm{max}}^{kin} = \frac{E_\mathrm{T}\, V_{\mathrm{max}}^{inv}}{V_{max}\, S_m} \quad \textbf{(13)}$$

**SI Derivation 3. Fraction of catalytically available sites $\boldsymbol{\Gamma_{\mathrm{max}}^{kin}}$.**[15,18]

**Langmuir Assumptions**

**1. Monolayer adsorption (one molecule per site)**

$$S + \mathrm{L} \rightleftharpoons SL$$

**2. Identical, energetically equivalent sites**

$$K_D = constant\ for\ all\ sites$$

**3. No interactions between adsorbed molecules**

$$\Delta G_{ads} = constant$$

$$K_D\ independent\ of\ \theta$$

**4. Fixed amounts of surface sites**

$$[S]_{tot} = [S] + [SL]$$

$$0 \leq \theta \leq 1$$

$$\theta = \frac{[SL]}{[S]_{tot}} = \frac{q_e}{q_{max}}$$

**5. Dynamic equilibrium (reversible binding)**

$$k_a C_e [S] = k_d [SL]$$

$$K_D = \frac{k_d}{k_a}$$

**Binding Equilibrium**

$$S + L \underset{k_\mathrm{d}}{\overset{k_\mathrm{a}}{\rightleftharpoons}} SL$$

$$K_D = \frac{k_d}{k_a} = \frac{[S][L]}{[SL]}$$

In aqueous solutions, $[\mathrm{L}] = \mathrm{C_e}$

$$K_D = \frac{[S][L]}{[SL]} = \frac{[S][C_e]}{[SL]}$$

$$\frac{[SL]}{[S]} = \frac{[C_e]}{[K_D]}$$

**Site Balance**

$$[\mathrm{S}]_{\mathrm{tot}} = [\mathrm{S}] + [\mathrm{SL}]$$

$$Let\ \theta = \frac{[SL]}{[S]_{tot}} = \frac{q_e}{q_{max}}$$

$$\frac{[SL]}{[S]} = \frac{\theta}{1-\theta}$$

Substitute into Binding Equilibrium

$$\frac{\theta}{1-\theta} = \frac{C_e}{K_D}$$

$$\theta = \frac{C_e}{K_D + C_e}$$

**Langmuir Isotherm**

$$\frac{q_e}{q_{max}} = \theta = \frac{C_e}{K_D + C_e}$$

Fractional site occupancy: $\theta = \frac{q_e}{q_{max}}$

$q_e$ =amount adsorbed at equilibrium

$q_{max}$ =maximum adsorption capacity

**SI Derivation 4. Langmuir binding isotherm.**[19–25]
